# Hygroregulation, a key ability for eusocial insects: Native Western European bees as a case study

**DOI:** 10.1101/351874

**Authors:** Iris Eouzan, Lionel Garnery, M. Alice Pinto, Damien Delalande, Cátia J. Neves, Francis Fabre, Jérôme Lesobre, Sylvie Houte, Andone Estonba, Iratxe Montes, Télesphore Sime-Ngando, David G. Biron

## Abstract

Sociality has brought many advantages to various hymenoptera species, including their ability of regulating physical factors in their nest (e.g., temperature). Although less studied, humidity is known to be important for egg, larval and pupal development. Two subspecies of *Apis mellifera* of the M branch, also called black bees, were used as models to test the “hygroregulation hypothesis”, by means of monitoring hygrometry in hives during one year in four conservation centers: two in France (*A. m. mellifera*) and two in Portugal (*A. m. iberiensis*). We investigated the ability of both subspecies to regulate the hygrometry daily, but also during the seasons and one complete year. Our data and statistical analysis probed the capacity of the bees to regulate humidity in their hive, regardless of the day, season or subspecies. Furthermore, the study showed that humidity in beehives is very stable even during winter, when brood is absent, and when temperature is known to be less stable in the beehives. These results would support that hygrometry could prevail over temperature in maintaining nest homeostasis, maybe because of a bigger importance of hygrometry for all the population during a year, or because of the ‘imprint’ of the evolutionary history of this hymenopteran lineage.

## Introduction

Terrestrial insects have high sensitivity to temperature and humidity. Most of them cannot control their body temperature (i.e., ectotherm species) and are, thereby, very dependent to their environment [1–3]. Indeed, temperature has an important impact on foraging [1] and it is also crucial for reproduction, larval and pupal development, and the success of the offspring [4]. Eusocial insects have also evolved many strategies to maintain their nest temperature stable and controlled [5,6], especially in order to protect the eggs from extreme temperature variations [6]. Nest homeostasis provides not only an incubator for the brood, but also a thermal refuge for individuals to use temperature gradients to regulate their own body temperature [7].

The eusocial bees use the thermoregulation to ensure their survival and health [3,6,8]; for instance *A. mellifera* larvae can only survive in an environment with slight temperature fluctuation (i.e., from 32 to 36 °C) [9]. The nest thermoregulation strategies of eusocial insects can be passive (e.g., nest orientation, architecture) or active (e.g., clustering, incubation), depending on the species [6]. Those thermoregulation strategies could have an impact on hygrometry levels [10], a less studied phenomenon, but also vital for terrestrial insects [11]. Indeed, environmental humidity is an essential factor to control for the survival of adults and eggs [11].While eusocial insects exhibit a range of humidity preferences in the nest regarding their activity or the presence of brood [12], they generally require a relative humidity (RH) of above 55% to hatch successfully, with the highest survival between 90 and 95% RH [4].

Here, we chose the honeybee, *Apis mellifera* (Linnaeus 1758) (Hymenoptera, Apidae), as model organism. This insect is worldwide present and it has been increasingly studied since the decline of its colonies in many occidental countries, due to its key role in pollination of many crops and in sustaining the wild biodiversity of ecosystems [13]. While several experiments have been conducted (i) to disentangle the mechanisms that lead to creation and maintenance of microclimate in a beehive [9,14], and (ii) to determine the impact of parasitism on thermoregulating social behavior [8], little is known about hygrometry in beehives. As for temperature, the eggs are very sensitive to humidity fluctuations within the nest [11]: a dry atmosphere can lead to the eggs death, either because the embryos die or because the dry egg envelops become too hard for the larvae to hatch [11]. After hatching, humidity is still essential for young bees, and considered at least as important as temperature for larval and pupal development [4,15,16].

In this study, we investigated for the first time humidity for one year in beehives of native European honeybee subspecies belonging to the M evolutionary branch (*mellifera* subspecies of Western Europe, also called black bees) [17]. Two subspecies of M lineage were studied in two European countries: *A. m. mellifera* in France, and *A. m iberiensis* in Portugal. Our experimental setup includes daily, seasonal and annual measures of hygrometry in the beehives with the main objective of assessing the capacity of bees to hygroregulate their nests (i.e. “hygroregulation hypothesis”), and in order to determine if geographic, seasonal or subspecies parameters could impact this capacity.

## Materials and Methods

### European study sites

The study was conducted in four conservation centers created to preserve *A. m. mellifera* in France and *A. m. iberiensis* in Portugal. These conservation centers, of an approximate extension of 350km^2^, are organized in a sanctuary zone encircled by a buffer zone, designed to prevent queens from mating with males from outside the conservatory area. The French conservation centers are located at Rochefort (48°35‘47’N; 1°57‘57’E) and Pontaumur (45°51’51”N; 2°40’24”E), in the regions of “Ile-de-France” and “Auvergne-Rhône-Alpes”, respectively. The landscape in Rochefort corresponds to that of “plain beekeeping”, and the landscape in Pontaumur corresponds to that of “semi-mountain beekeeping”. The Portuguese conservation centers are located at two latitudinal extremes in Gimonde (41°48’31”N; 6°42’41”W) and Zavial (37°03’14”N; 8°52’40”W), in the regions of “Trás-os-Montes” and “Algarve”, respectively. The landscape in Gimonde corresponds to that of “semi-mountain beekeeping”, and the landscape in Zavial corresponds to that of “plain beekeeping”.

### Experimental design to monitor the hygrometry in hives

In each conservation center, six healthy beehives from the sanctuary zone were randomly chosen to be monitored. relative humidity (RH) was measured by three thermo-hygro button data loggers (also named iButtons). The iButtons were placed in the nest of each of the six monitored colonies, at the two outermost brood frames (iButton A and C) and at the central brood frame (iButton B), each of them hanging between two frames by an iron thread. An external thermo-hygro button was placed in each apiary at approximately 200-cm high, near the monitored beehives, to register environmental RH variation. Each iButton was programmed to measure RH hourly with a precision of ±1%. Data from iButtons were collected every four months by a reader (Plug & Track, Progues PLUS) and exported to Excel format using the Thermotrack PC V.7 software. The pluviometry data was daily collected at 2 a.m., 8 a.m., 2 p.m. and 8 p.m. by a weather station (Micro El d.o.o., Zagreb) placed in each sanctuary.

### Statistical Analysis

To test the “hygroregulation hypothesis” at daily scale, first, RH (%) measured for each hive was plot to give an overview of the humidity evolution within the hives. Then, the days with the most similar external RH daily profile between the four geographic locations and for two contrasting seasons in beekeeping: summer (high colony activity) and winter (low colony activity) were selected using a similarity index (S) for days without rain in the four conservatories. A similarity matrix S=1−D, where S denotes similarity and D denotes distance (D (x,y) = (∑_i_ (x_i_ − y_i_)^2^)^1/2^), with x_i_ and y_i_ being the peers of conservatories to compare, was calculated using Statistica 8.0 software (Stat Soft Inc., Arizona, USA). This index allowed to select one day in winter and on day in summer to compare in-hive RH and external RH in each conservatory. Then, both days were divided into two parts: the part of the day with gradual decrease in external RH (i.e., from maximum RH, at 6 a.m., to minimum RH, at 5 p.m.), and the part of the day with a gradual increase of external RH (i.e., from the minimum RH of the day, at 6 p.m., until next day’s morning, when RH was maximum, at 5 a.m.). For each selected day, linear regression was calculated using in-hive from ibutton B and external data, for each day part, to test the daily “hygroregulation hypothesis” (i.e. weak positive correlation (0 < r ≤ 0,6) linked with a R^2^ ≤ 0,40 (weak coefficient of determination), or negative or null correlation (r ≤ 0). An index ((RHi-RHe)/RHe) with RHi being the in-hive data of iButton B and RHe being the external RH, was calculated for each hour of each selected day. A clustering analysis was performed with this index in order to classify the hives from each conservatory in a hierarchical way. This analysis was performed using the PermutMatrix 1.9.4 seriation software (SupAgro, Montpellier, France).

An analysis of RH was also performed for one complete year, from September 2015 to August 2016. In-hive data from all iButtons and beehives were put together for each conservation center to obtain a global mean and standard error and compare them with the external data, using Statistica 8.0. A statistical analysis was also conducted at the season level, by considering September to November for autumn, December to February for winter, March to May for spring and June to August for summer.

## Results

### Daily hygroregulation

The external RH from the two selected days for the four conservation centers are given in S1 Fig: December 11th, 2015, a winter day without rain or snow, had the most similar external RH profile among the four conservation centers (50% similarity), whereas July 21st, 2016, was the most similar not-rainy summery day (40% similarity). Fig 1 shows the regression lines calculated for July 21, 2016, for the central iButton (B) of hive 1 in each conservation center. These are representative of the regression lines for the remaining five hives and iButtons (i.e., linear equations, r and R2), that are shown in supplementary material S1 Table. The cluster analysis was only performed for iButton B because it assessed RH in the main part of the brood (i.e. eggs, larvae and pupae) (Fig 2). During summer, the results show globally a positive regulation (in-hive RH > external RH) in the afternoon, especially in Gimonde. In Gimonde, the beehives are mostly maintained with low humidity inside, compared to the other apiaries. During winter, results show a similarity between all conservation centers, since RH is maintained along the whole day at lower level inside than outside the hives (S3 Fig).

**Fig 1.**
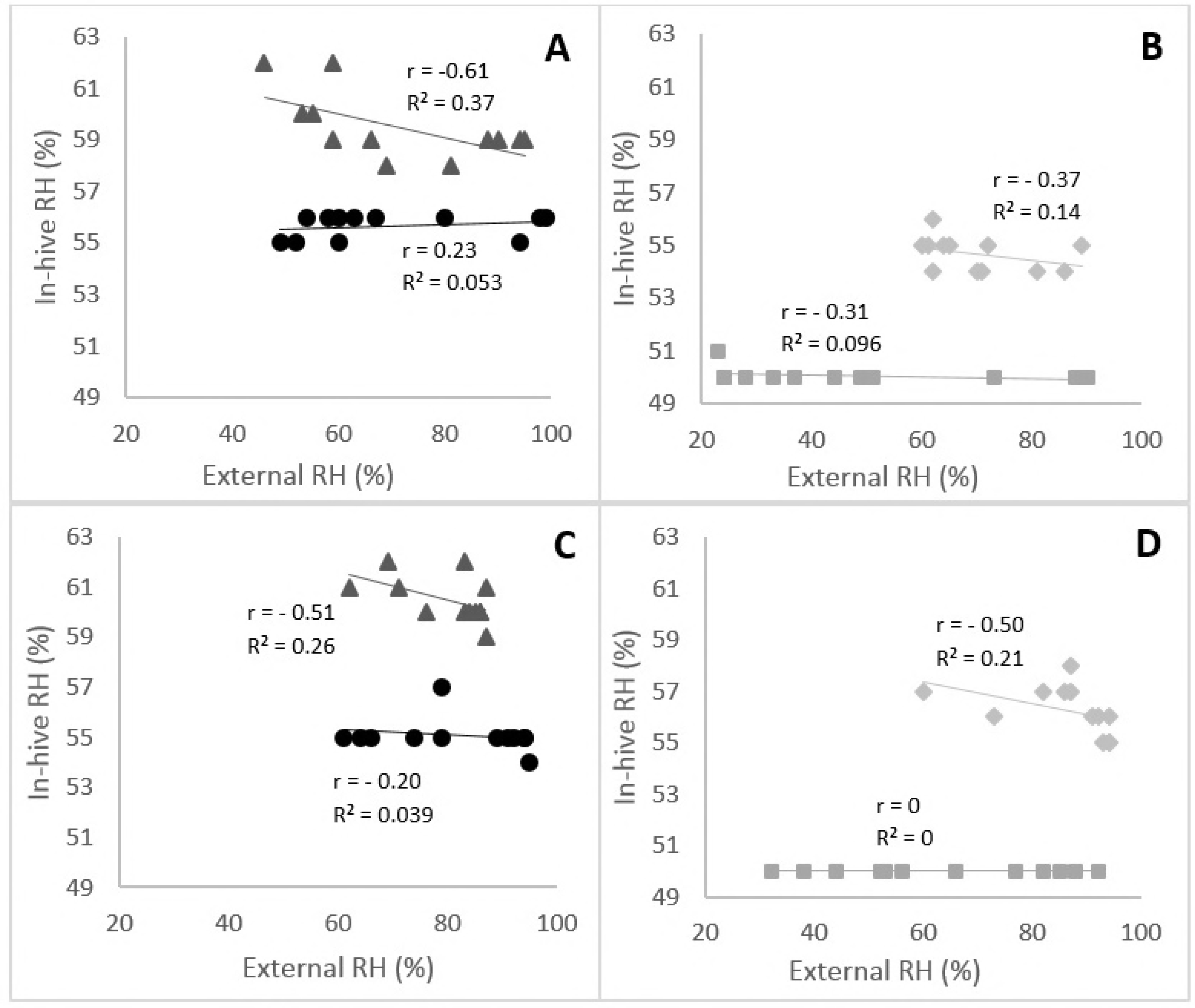
Linear modeling of the relationship between RH observed in the summer in the beehives 1, iButton B (in-hive) with those recorded in their habitat (external) in each conservation center. The modeling was done for one day (July 21, 2016) and separated in two parts: downward RH from 6 a.m. to 5 p.m. and upward RH from 6 p.m. to 5 a.m. the next day. Linear modeling for *A. m. mellifera* is represented in A (downward RH) and C (upward RH) for Pontaumur (▴) and Rochefort (•). Linear modeling for *A. m. iberiensis* is represented in B (downward RH) and D (upward RH) for Zavial (♦) and Gimonde (▪).

### Seasonal and annual hygroregulation

Average annual in-hive RHs were significantly different from average annual external RHs (Mann-Whitney, p < 0.05) for the two honeybee subspecies deployed in the four conservation centers (*A. m. mellifera* in Rochefort and Pontaumur, *A. m. iberiensis* in Gimonde and Zavial) (Fig 3). In contrast, the comparison of the average in-hive RH shows that, while Gimonde is similar to Pontaumur, the other conservation centers are statistically different from each other (Multiple Comparison, p < 0.05).

**Fig 3.**
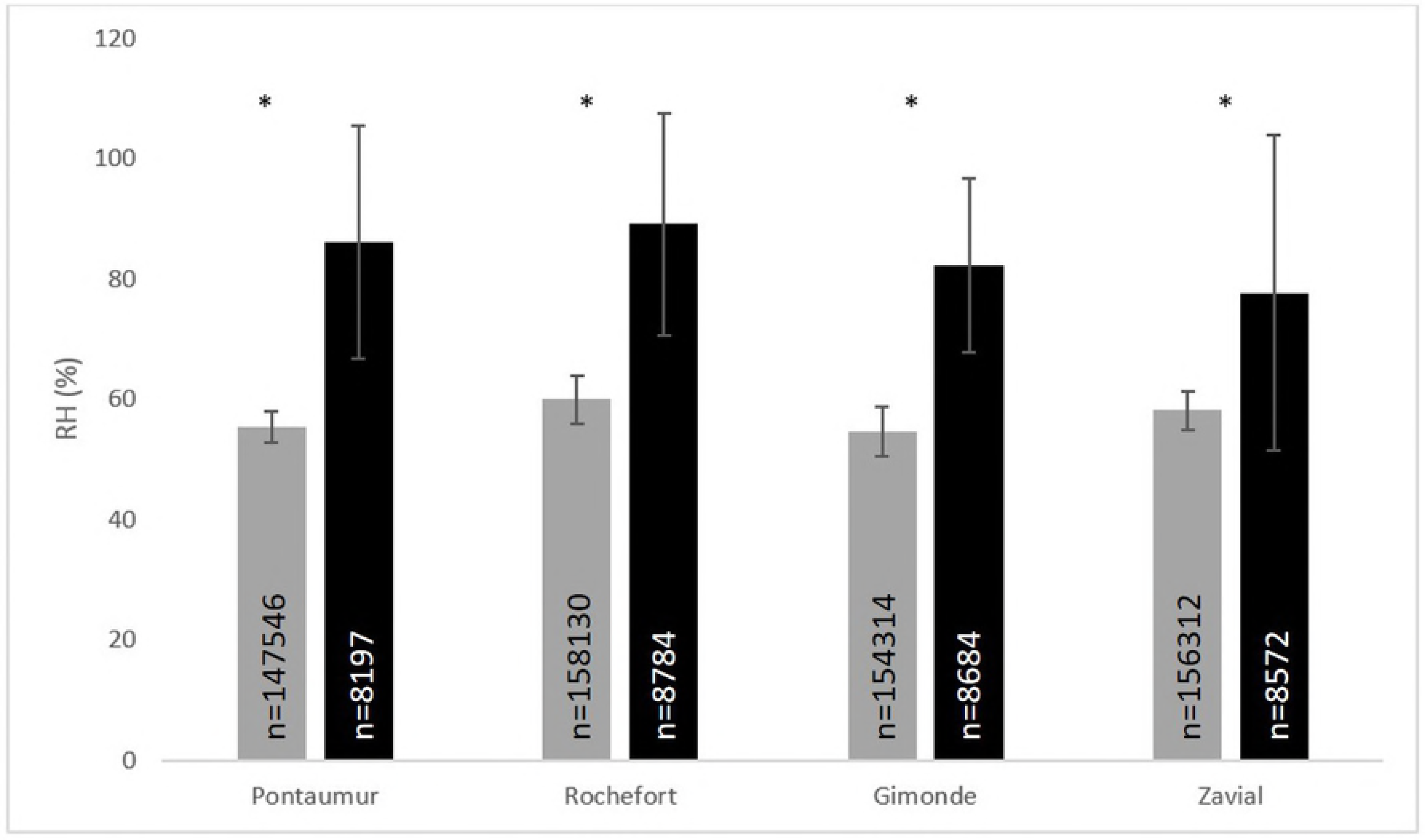
Mean and standard errors of RH levels over a complete year (September 2015 - October 2016) for the four conservation centers. In-hive RH (grey) were calculated using the mean of the three iButtons (A, B and C) and the six beehives for each conservation center, and external RH (black) was calculated using only the external iButton for each conservation center. “*” = in-hive and external RH are different (Multiple Comparisons, p<0.05). Detailed mean and standard errors for the four seasons are shown in supplementary material Fig S4.

The seasonal analysis of RH showed significantly differences of RH between the in-hive and the external data (Mann-Whitney, p < 0.05), for each conservation center in each season (supplementary material, Fig S4). The average RH inside the beehives was maximum at 61.0 ± 1.3% (Zavial, winter) and minimum at 50.4 ± 3.0% (Gimonde, autumn). Furthermore, the statistical analysis shows that, for each season, the four conservation centers had significantly different in-hive RH (Multiple Comparisons, p < 0.05), except in spring, when Rochefort (60.2 ± 3.2%) was similar to Zavial (59.5 ± 2.0%) (Multiple Comparison, p = 1.00).

## Discussion

Eusociality brought advantages to many insect species, especially the ability of living in community and, thus, to regulate some parameters of their immediate environment, such as temperature and humidity. While temperature regulation in eusocial insects has been studied extensively [1,5,6,14,18] providing insights about the factors that influences temperature management [1,19,20], such as climate [21] or genetics [18], there is little information on the humidity regulation inside nests and hives. In this study we tested the “hygroregulation hypothesis” by means of using as model two native honeybee subspecies from the M branch, *A. m. mellifera* and *A. m. iberiensis,* and the assessment of the temporal hygrometry variation inside and outside beehives. Thereby, we sought to better understand the ability of eusocial insects to regulate the humidity of their nest (hive).

Our study has the advantage to present an approach at different scales, namely the day, the season and the year, in geographic regions with contrasting climates and landscapes. Our results overall show low or even no correlation between humidity inside and outside the hive for the two honeybee subspecies, in summer (Fig 1, supplementary material S1 Table) and in winter (S2 Fig, S1 Table). Thus, the black honeybee populations in French and in Portuguese conservation centers maintain the nest humidity constant over the day, despite the extreme variations that take place outside (Fig 2). In addition, the results show the ability of the two honeybee subspecies to regulate their nest humidity regardless the season, even in the heart of the summer or in the winter, when brood is absent [22]. Furthermore, no differences were found over the year between the two honeybee subspecies, since the in-hive hygrometry in Pontaumur (*A. m. mellifera*) and Gimonde (*A. m. iberiensis*) are at the same level, and those from Zavial and Rochefort are higher and close to each other, with Rochefort having always a higher hygrometry level (58.1 ± 3.2% for Zavial and 59.9 ± 4.1% for Rochefort over the year). The similar in-hive hygroregulation in Pontaumur and Gimonde on the one hand, and of Zavial and Rochefort on the other hand, could be due to the environment, Pontaumur and Gimonde being in a semi-mountain place, and Rochefort and Zavial being both in a plain landscape.

**Fig 2.**
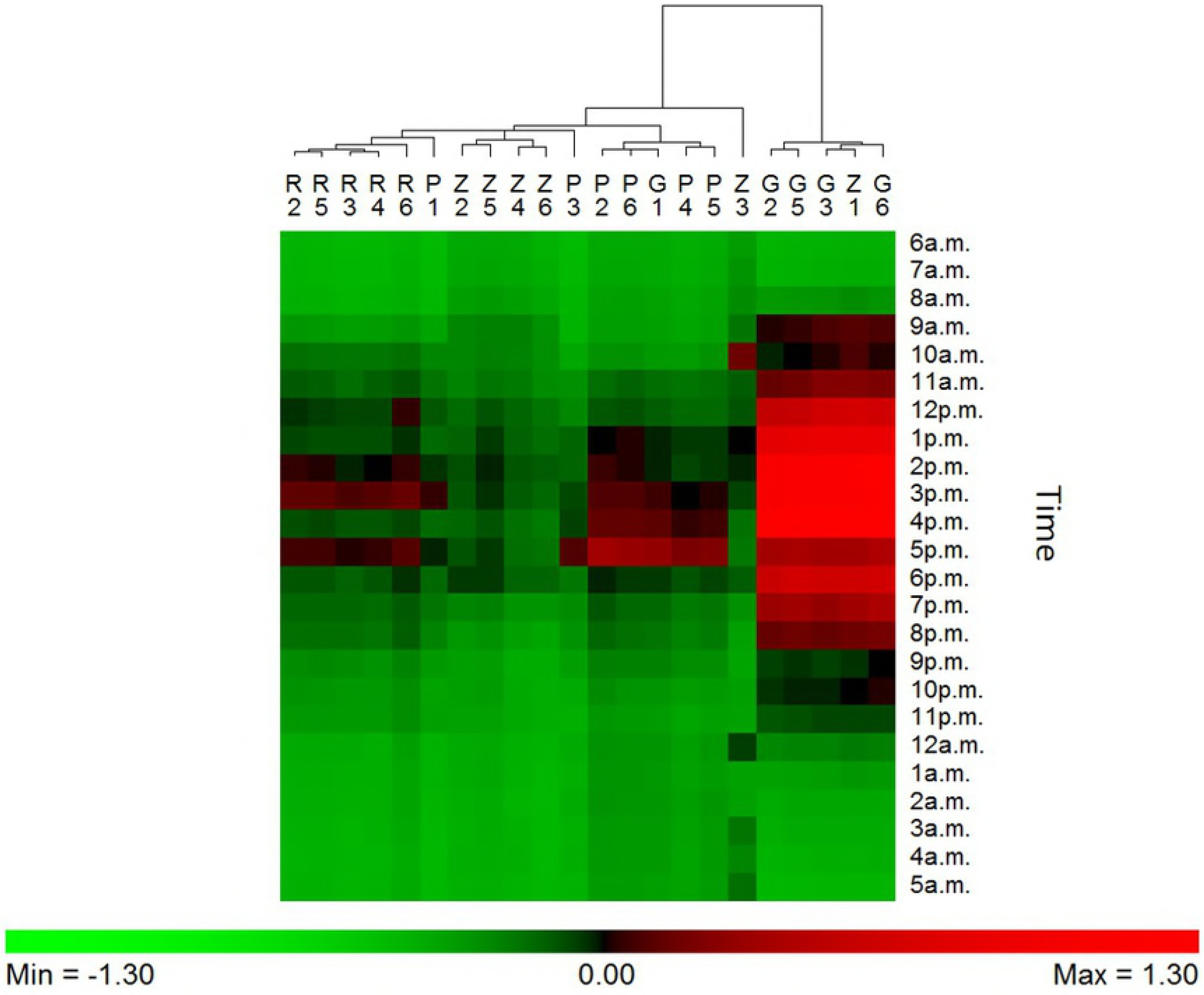
Hygroregulation observed in the central brood frame (iButton B) of each of the 6 hives during a 24h-period in the summer, from 6 a.m. (July 21, 2016) to 5 a.m. of the next day (July 22, 2016), in the four conservation centers (Rochefort, R; Pontaumur, P; Gimonde, G; and Zavial, Z). Green means that the in-hive RH is lower than the external RH (negative regulation), red means the opposite (positive regulation).

It has been shown that temperature is better regulated in the beehives in the summer than in the winter, when there is brood in the hive, [23]. This is likely due to the important role of temperature in egg development; it is known that at too low (i.e. < 33°C) or too high (i.e. > 36°C) temperature eggs, larvae and pupae die [9,24]. Similarly, maintenance of the humidity level is crucial for social insects; it directly affects the proper development of eggs, larvae and pupae [4,11], which die when the ambient environment is too dry [11]. Therefore, a minimum of 55% humidity is required for honeybee eggs to hatch, with a maximum survival rate between 90 and 95% [4]. In addition, Ellis *et al.* (2008) have shown that honeybee workers have a marked preference for approximately 75% humidity in the absence of brood [16]. In this study, we obtained lower humidity levels in the beehives, with seasonal RH ranging from 50.4 ± 4.4% (Gimonde, autumn) to 65.2 ± 7.8% (Rochefort, winter) (supplemental material Fig S4). Seasons when brood is present in the hive (spring and summer) do not differ significantly from the rest of the year (supplemental material Fig S4). However, in those two seasons, RH does not drop below 50%, threshold below which eggs cannot hatch [4].

The fact that humidity is stable and, therefore, regulated in the winter, unlike temperature [14,23] suggests two hypotheses: (i) the humidity is more easily maintained in the hive than the temperature, despite the fact that the two honeybee subspecies must reduce their energy efforts because food reserves are limited [25], and (ii) moisture is a more important factor for adult health than previously suggested in the literature [4,11,15,16], especially for adult bees that keep it constant even in the absence of brood. The first hypothesis involves a link between temperature and humidity. Indeed, honeybee subspecies, such as *A. m. mellifera* and *A. m. iberiensis*, mainly use active regulation systems to manage the temperature of their nest [6]. This strategy includes ventilation, which has also an impact on ambient humidity [22,23]. The second hypothesis is based on studies by Buxton, who showed in 1932 that insects only drink very rarely and need a moist environment to avoid desiccation, whether they are in the larval or adult stages [11,16].

It is important to note that the evolutionary history of honeybees goes back several million years [17], whereas their encounter with humans dates back only 15,000 years ago [22], when bees moved from nesting in various natural cavities to beehives. Their ability to maintain stable moisture and temperature within the colony may have facilitated the migration of different subspecies to geographical areas with a climate that often varies greatly with the seasons, or particularly arid countries, such as in many parts of Africa [26].

Currently, global warming is causing significant changes in the environment that organisms have to face [27–31]. Since 1990, the average global surface temperature, the environmental factor with the greatest impact on the biosphere, has increased by around 0.9°C, with a faster rise for the minimum than for the maximum [31]. This global warming contributes to the destruction of several habitats and biological invasions in several ecosystems [30,31]. Insects, however, show strong adaptabilities to new climates, for example by modifying their range [30,32] or their period of activity [33]. However, according to our results, the two honeybee subspecies included in this study require a nest with relatively stable and high humidity levels. Some eusocial hymenoptera living in relatively arid areas have adapted their behavior according to their unavailability of water [2,33]. For those social species, the lack of water due to global warming could lead to a significant change in their geographic distribution, in order to survive to those new and limiting conditions.

Moreover, as our data suggest a high importance of moisture for both brood and adults, it is conceivable that the development of certain diseases may be manifested by a disturbance of the humidity in the nest: either (i) poor moisture control that would favor the occurrence of opportunistic parasites such as varroa mites whose ability to reproduce is impacted by moisture in hives [34], or (ii) the presence of parasites and pathogens causing weakening of colonies [35–37], which would induce a decrease in the ability of insects to properly regulate the humidity of their nest. Thus, we suggest that monitoring abiotic factors such as humidity and temperature in honeybee hives could be a strategy for identifying colonies having disturbance in their normal functioning as a eusocial community, and help to find the eventual factors leading to the decline of a honeybee colony.

## Conclusion

Our data and statistical analysis sustain the validation of the “hygroregulation hypothesis”: the eusocial honeybee ability to regulate the humidity of the hive (nest) at a day, but also at seasonal and year scales. Thereby, humidity is constant during the year in the beehives, even in winter when temperature is less regulated because of the absence of brood. Furthermore, no differences were showed between the two black bee studied subspecies, *A. m. mellifera* and *A. m. iberiensis*. However, regardless of the genetics, it seems that the landscape could have a bigger impact on the regulation, the apiaries in the plain having an in-hive hygrometry slightly lower than those in the semi-mountain landscape. Overall, our results help to better understanding how hygrometry is regulated in eusocial insects, and its relative importance compared to temperature.

## Acknowledgements

We thank Noel Mallet, Claude Grenier, Jean-Charles Labat, Céline Robert, Paulo Ventura, Miguel Vilas-Boas, Jonathan Gaboulaud, Cécile Ribout, Jean-François Odoux, Egoitz Galarza and Hélène Legout, who all helped us in the BEEHOPE project.

## Supporting information

**S1 Table. Linear equation, r and R^2^ for each model of the relationship between RH observed in the summer (A) and in the winter (B) in the beehives (in-hive) with those recorded outside (external).** The empty places correspond to iButtons that were absent from the beehive at this moment. Missing data are due to a breakdown for two iButtons for a couple of days. **Figure S1:** Relative humidity
(RH) measured by the external iButton in the four conservation centers: Pontaumur (▴), Rochefort (•), Zavial (♦) and Gimonde (▪), (A) in summer (July 21, 2016) and (B) in winter (December 11, 2015). These two dates were chosen because they have the most similar external RH among the four conservatories in the two seasons, and because it did not rain in any of the four conservation centers. The data were taken from 6 a.m. to 5 a.m. the next day for both dates.

**S2 Fig. Linear modeling of the relationship between RH observed in the winter in the beehives (in-hive) with those recorded in their habitat (external).** The modeling was done for one day (December 11, 2015) and separated in two parts: downward RH from 6 a.m. to 5 p.m. and upward RH from 6 p.m. to 5 a.m. the next day. Linear modeling for *A. m. mellifera* is represented in A (downward RH) and C (upward RH) for Pontaumur (▲) and Rochefort (•). Linear modeling for *A. m. iberiensis* is represented in B (downward RH) and D (upward RH) for Zavial (♦) and Gimonde (▪).

**S3 Fig. RH regulations observed in winter for each colony (iButton B) regarding to the time of the day: from 6 a.m. to 5 p.m. the next morning.** Green means the in-hive RH is lower than the external hygrometry (negative regulation), red means the opposite (positive regulation).

**S4 Fig. Mean and standard errors of in-hive (grey) and external (black) RH for autumn (September-November, 2015), winter (December 2015-February 2016), spring (March-May, 2016), and summer (June-August, 2016).** The mean and standard errors for in-hive data were calculated over the 3 iButtons (A, B and C) for the six beehives of the four conservation centers and external data were calculated using over the external iButton. “*” = in-hive and external RH are different (Multiple Comparisons, p<0.05).

